# Intron architecture predicts chromatin features in *Arabidopsis thaliana*

**DOI:** 10.1101/2025.10.15.682614

**Authors:** Alice Pierce, Alan Rose, Grey Monroe

## Abstract

Introns are ubiquitous yet enigmatic features of eukaryotic genomes. They are reported to affect gene expression regulation but the mechanisms remain largely unresolved. Here we investigate the connections between intron architecture and chromatin features – histone marks, histone variants, DNA methylation, and gene expression patterns in *Arabidopsis thaliana*. We found that first intron positions predict active chromatin marks found near transcription start sites while the number of introns explains different chromatin marks enriched in the core of gene bodies in broadly expressed genes. Notably, gene body chromatin marks exhibit a positional gradient distribution across the ordinal position of exons and introns. We tested these relationships by comparing recent gene duplicates with diverged intron architectures, and confirmed that intron number is positively associated with H3K4me1, H3K36me3, H2A.X, meCG, and broad expression across tissues. Our results suggest two distinct mechanisms in which plant intron architecture may affect chromatin states and ultimately gene expression, motivating future experiments: intron positions early in genes may affect the establishment of activating histone marks around transcription start sites, while a greater total number of introns may increase the number of intronic motifs and gene length in general, allowing for the increased accumulation of gene-body-associated chromatin features.

**Short summary:** Here we investigate the association between intron architecture and chromatin features in *Arabidopsis thaliana*. We found that the position of the first intron is negatively correlated with marks found near the transcription start site and associated with active expression. We also found that the number of introns within genes is correlated with marks found in gene bodies and are associated with actively transcribed genes.

## Introduction

Introns, non-coding regions of DNA that are interspersed within genes, are ubiquitous components of eukaryotic genomes. The discovery of introns in 1977 was a major breakthrough in molecular biology, highlighting the essential role of introns in gene structure and expression (Berget et al. 1977). Research has since demonstrated that introns contribute to gene expression regulation through various mechanisms. Some examples in which introns affect gene expression include influencing transcript stability (Duan et al. 2013; Narsai et al. 2007; Sharova et al. 2009; Wang et al. 2007), mRNA export (Valencia et al. 2008), and harboring cis-regulatory elements (Voichek et al. 2024). These findings have provided insight into the functional complexity of intronic sequences and their role in eukaryotic gene regulation.

Further adding to the complex nature of intron biology is the less-understood phenomenon of Intron-Mediated Enhancement (IME). IME introns can boost gene expression; however, the mechanism by which they regulate gene expression remains unclear in plants. In yeast, IME is thought to work in a splicing-dependent manner, altering the 3D architecture of the genes to enhance gene expression (Moabbi et al. 2012; Dwyer et al. 2021). In plants, however, the mechanism by which stimulatory introns are capable of boosting expression remains unclear, but experiments suggest that a subset of introns boost expression through a splicing-independent mechanism (Rose and Beliakoff 2000; Rose 2002). Recent hypotheses suggest that stimulatory introns may have a role in establishing favorable chromatin states of their genes, facilitating the deposition of epigenomic marks such as H3K4me3 and H3K9ac (Gallegos and Rose 2015) as previous work has shown that the distance between the transcription start site (TSS) and the first intron to be associated with H3K4me3, H3K9ac and gene expression(Bieberstein et al. 2012; Girardini et al. 2023). These marks are typically deposited near the TSS and are associated with actively transcribed genes (Leng et al. 2020; Zhang et al. 2009; Charron et al. 2009). However, further work is needed to determine which marks, if any, are associated with introns, and what features of introns are important to establish these relationships.

To investigate relationships between intron architecture and chromatin features, we reanalyzed a suite of previously published ChIP-seq, ATAC-seq (Liu et al. 2018), cytosine methylation (Monroe et al. 2022), and gene expression data (Mergner et al. 2020) in *Arabidopsis thaliana* (*Arabidopsis*, hereafter). We demonstrate that intron features are correlated with chromatin features that have consequences on the expression of their genes. Furthermore, we used gene duplicates to infer the causality of the genome-wide patterns observed by comparing the abundance of chromatin marks on gene duplicate pairs that have diverged in intron content.

## Materials and Methods

### Gene Feature Annotation

The *Arabidopsis thaliana* TAIR10 reference genome (Lamesch et al. 2012) was used to obtain gene, exon, and intron coordinates. The dataset was filtered to include only representative gene models. The number of introns, gene length, intron length, exon length, intron percent, first intron length, and first intron position relative to the transcription start site were calculated using custom R scripts.

### Chromatin Features

#### Epigenomic data

Histone variants, post-translational modifications, and chromatin accessibility datasets were accessed from a collection of ChIP-seq and ATAC-seq datasets in BigWig format maintained by the Plant Chromatin State Database (Liu et al. 2018). These included data on 15 histone post-translational modifications, 5 histone variants, 5 DNA-associated proteins, and chromatin accessibility (**Supplementary Table S1**). The read depth of chromatin features was calculated in 10-bp windows (no overlap) across the genome. The scaled mean across all datasets for each chromatin mark was calculated and used for downstream analyses.

#### Cytosine methylation data

Previously published DNA cytosine methylation datasets were obtained (Monroe et al. 2022; Kawakatsu et al. 2016). The number of methylated sites for each methylation context (CG, CHH, CHG) was calculated across gene bodies and normalized by the total number of cytosines found across gene bodies, exon sequences, or intron sequences.

#### Tissue-specific gene expression

Tissue-specific expression from 54 tissues was obtained from previously published data (Mergner et al. 2020). Missing values were set to 0. The expression datasets were averaged to obtain the mean expression, from which we also extracted the coefficient of variation of the expression across tissues for each gene. Tissue breadth was calculated by counting the number of tissues in which the expression is non-zero.

### Genome-wide Analyses

#### Correlation

The Spearman correlation between gene features (Gene Length, Total Exon Length, Number of Introns) and chromatin features was calculated for all genes annotated in the TAIR10 genome. The Spearman correlation between intron features (Total Intron Length, Intron Percent, First Intron Position relative to the transcription start Site, and First Intron Length) and chromatin features was calculated for intron-containing genes (**Supplementary Table S2, Supplementary Table S3**).

#### Random Forest Regressor

##### Random Forest models

We employed random forest regressor models using the R package “randomForest” for each chromatin feature across intron-containing genes, using gene features (Gene Length, Total Exon Length, Number of Introns, Total Intron Length, Intron Percent, and First Intron Length) as the predictor variables. Consistent with prior work, to minimize the effect of outliers, we ranked the response variables and resolved ties using the average value. For each model, we calculated the percent variance explained in the response variable explained by the predictor variables.

##### Random Forest model evaluation

To test the performance of each model, 5-fold cross-validation was performed on an 80-20 train-test dataset. From the cross-validation, we extracted the RMSE, MAE, R^2^, and Pearson Correlation between actual and predicted values from the models (**Supplementary Figure 2, Supplementary Table S4**). Feature importance for each model was calculated as the percent increase in mean square error after permuting each predictor variable, averaged across 5-fold cross-validation (**Supplementary Table S5**).

#### Metaplots

We binned intron-containing genes by first intron position relative to the transcription start site, and number of introns, and plotted the mean distribution of the scaled read depth per group across 10 base pair windows, either 3kb or 1.5kb upstream and downstream from the transcription start site.

### Gene duplicates analysis

#### Datasets

Datasets of gene duplicates and transposed genes previously published were obtained (Kenchanmane Raju et al. 2023). The number of introns for each gene was calculated using custom R scripts. Epigenomic data was obtained from the Plant Chromatin State Database (Liu et al. 2018) and analyzed as described above.

#### Gene duplicates analysis

We calculated the difference in the mean scaled read depth (analysis described above) between gene duplicate pairs with no difference in intron number (n=6203), differ in 1 intron (n=2461), differ in 2 or 3 introns (n=959), or differ in more than 3 introns (n=366). We performed the Welch Two Sample t-test for the difference in means of read depth between duplicates with less and more number of introns.

#### Transposed duplicates analysis

We calculated the difference in the mean scaled read depth (analysis described above) between parental and transposed duplicates with no difference in intron number (n=1109), loss of introns (n=497), or gain of introns (n=356). We performed the Welch Two Sample t-test for the difference in means of read depth between transposed duplicates that lost introns and gained introns.

## Results

### Intron architecture is correlated with chromatin features commonly found near the TSS and throughout the gene body of actively transcribed genes

To establish if introns have possible effect on chromatin structure, we first aimed to assess whether univariate descriptions of intron architecture (first intron position, first intron length, intron number, total intron length, and intron percent) are correlated with chromatin features. We analyzed publicly available data maintained by the Plant Chromatin State Database for *Arabidopsis (Liu et al. 2018)*. Additionally, we analyzed previously published DNA methylation data (Monroe et al. 2022) and tissue-specific gene expression data (Mergner et al. 2020). The Spearman Correlation was calculated to evaluate the relationship between gene features and chromatin features. To further explore feature importance and to take into account the non-linearity of the biological data, we implemented random forest models for each chromatin feature, using gene features as predictor variables and employing 5-fold cross-validation with an 80-20 train-test split for robust evaluation.

The results showed that intron architecture is correlated with chromatin features (**Fig. 1, Fig. S1**). Notably, the position of the first intron relative to the TSS is positively correlated with histone post-translational modifications typically located near the TSS, such as H3K4me3 (rho=0.253), H3K4me2 (rho=0.267), H3K23ac (rho=0.234), and H3K14ac (rho=0.195) (**Supplementary Table S2**). Our random forest models also associated the first intron position as a key feature to predict these chromatin marks (**Fig. 1B**). In addition, we observed that the number of introns in genes is correlated with chromatin features typically found throughout gene bodies. These included features such as H3K4me1 (rho=0.514), H2A.X (rho=0.517), H3K36me3 (rho=0.4), and H3K27me3 (rho=-0.273) (**Supplementary Table S2**). Random forest analysis also highlighted intron number as an important predictive feature for these gene body-associated marks (**Fig. 1B**). These findings suggest that both the position and number of introns contribute to the regulation of chromatin structure in *Arabidopsis*.

**Figure 1.**
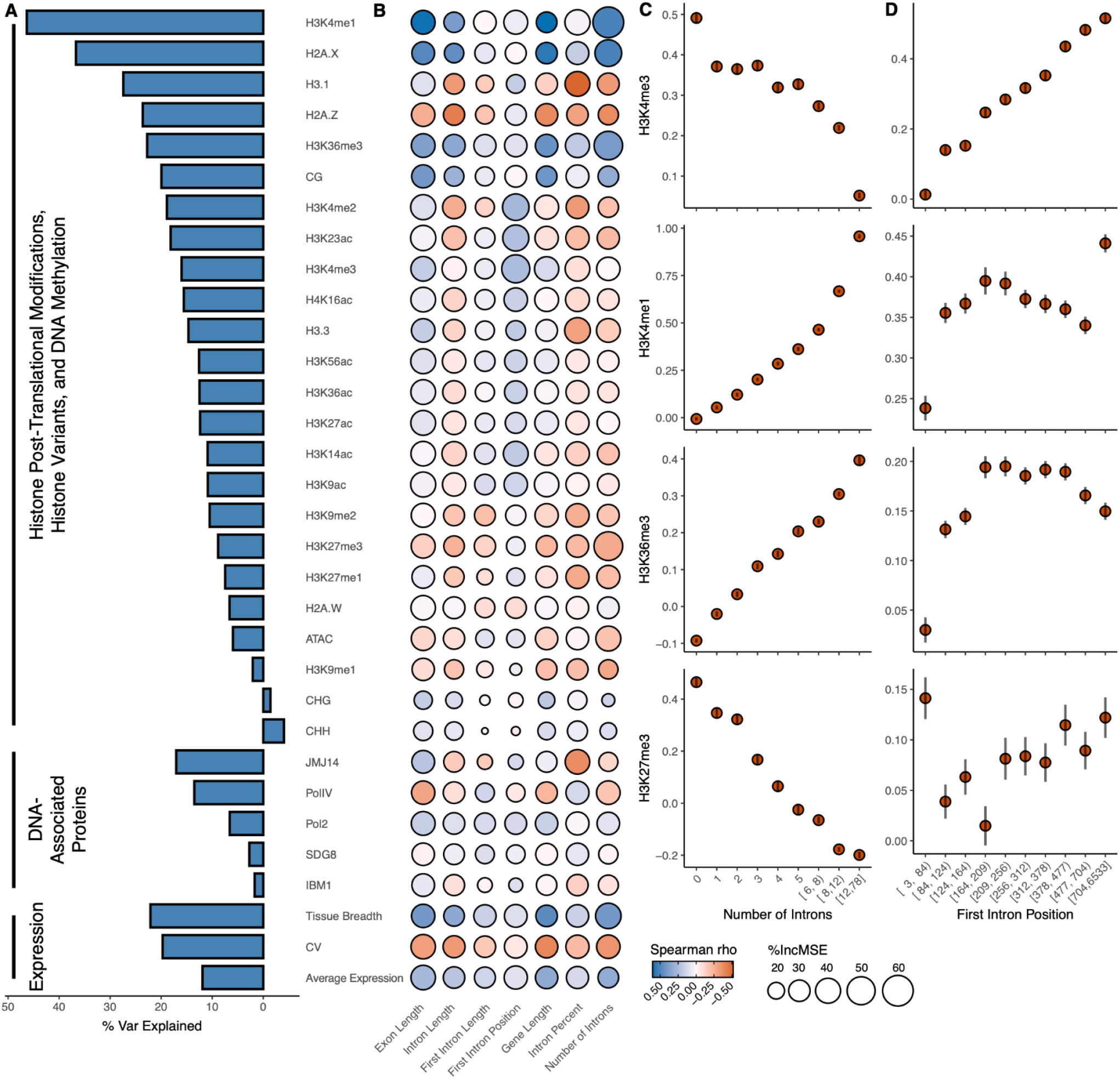
Prediction of chromatin features from intron architecture. **(A)**Bar graph showing the percent variance explained from random forest regression between gene features predicting each chromatin feature, DNA-associated protein, or gene expression feature across the gene body. **(B)** Correlation heatmap between gene architecture, chromatin features, DNA-associated features, or gene expression features across gene bodies. The color indicates the Spearman correlation coefficient between gene features and each chromatin feature. Size indicates the percent increase in mean square error introduced in the random forest regressor when the corresponding predictor is permuted. **(C)** Scatter plots showing the relationship between Intron Number or **(D)** First Intron Position relative to the Transcription Start Site for H3K4me3, H3K4me1, H3K36me3, and H3K27me3. Error bars represent ± standard error. Intron Number bins: 0 = 5939; 1 = 4118; 2 = 3080; 3 = 2470; 4 = 2095; 5 = 1625; [6,8) = 2520; [8,12) = 2976; [12,78] = 2383. First Intron Position bins: N: [3,84) = 2177; [84,124) = 2086; [124,164) = 2122; [164,209) = 2148; [209,256) = 2107; [256,312) = 2127; [312,378) = 2136; [378,477) = 2115; [477,704) = 2129; [704,6533] = 2120.

### H3K36me3 and H3K4me1 enrichment follows gene body position rather than exon-intron identity

We sought to determine whether chromatin features correlated with intron architecture exhibit differential distribution between introns and exons. To investigate this, we grouped genes by their total number of introns and plotted the mean scaled depth of chromatin marks across ordinal number of introns and exons (**Fig. S3**).

We found that the distribution of gene body marks such as H3K36me3 and H3K4me1 can be explained by the architecture and positional context within the gene body, rather than by feature identity (i.e. exon or intron) (**Fig. 2A, Fig. S3**). For instance, although intron 1 has higher H3K4me1 levels than exon 1, this difference does not persist when comparing intron 1 to exon 2, indicating that enrichment of these chromatin marks follow a positional order in the gene body.

**Figure 2.**
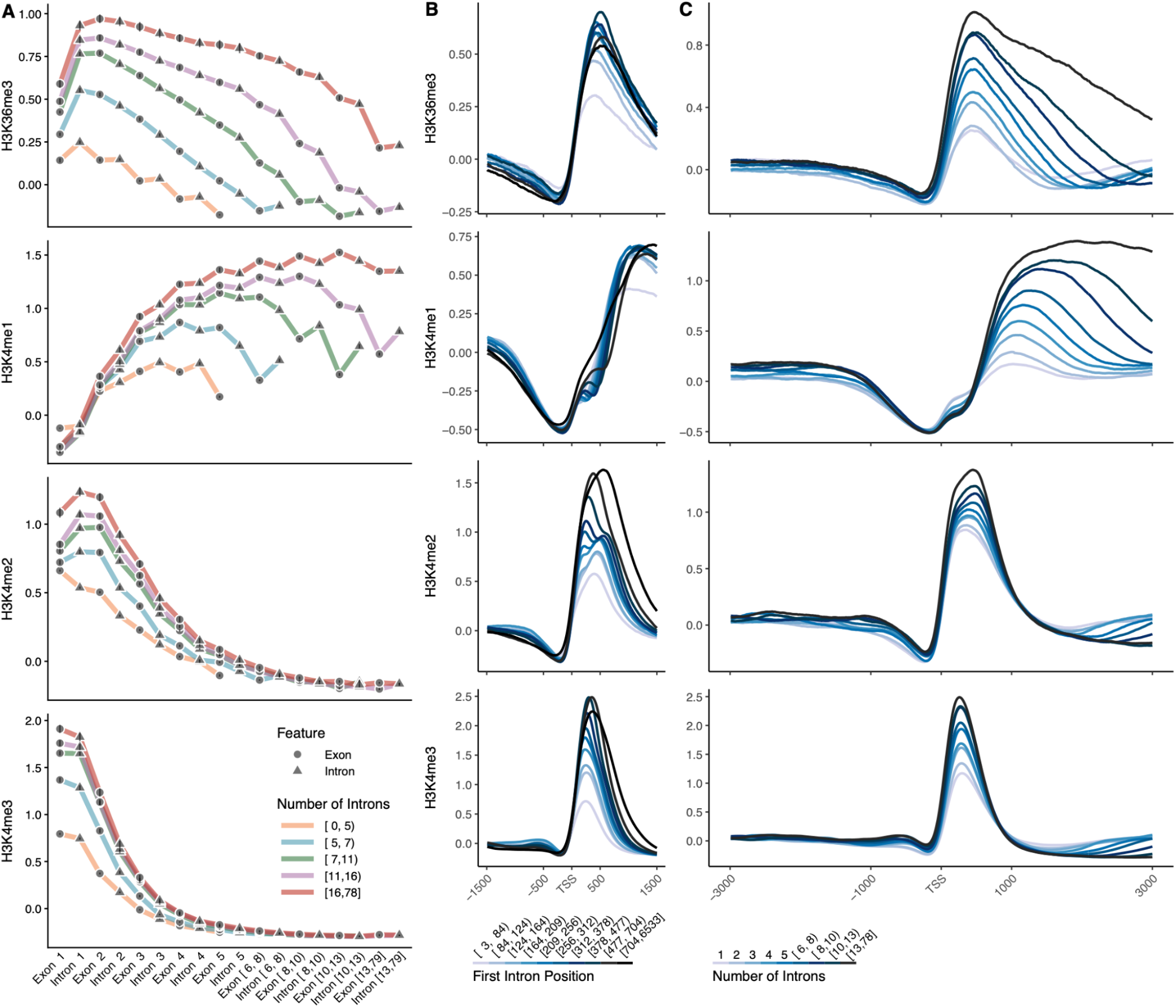
Relationship between intron features and histone post-translational modifications. **(A)**Scatter plots showing the mean scaled read depth of histone post-translational modifications across ordinal exons and introns. Genes were grouped by their number of introns. Circles correspond to exons and triangles correspond to introns. Error bars represent ± standard error. **(B)** Mean distribution of histone post-translational modifications of intron-containing genes grouped by first intron position relative to the TSS. **(C)** Mean distribution of histone post-translational modifications of intron-containing genes grouped by number of introns.

To further investigate the relationship between first intron position, number of introns, and chromatin features, we analyzed the average distribution of chromatin marks across gene bodies, binning genes by gene features (**Fig. 2B**). We found that for chromatin marks typically enriched near the TSS of actively transcribed genes, such as H3K4me3, a greater distance of the first intron from the TSS corresponds to higher average “peak” or amplitude in the distribution of these marks near the TSS (**Fig. 2B**).

For chromatin marks distributed throughout the gene body, such as H3K4me1 and H3K36me3, our analysis showed that more introns in genes is correlated with a higher abundance of these marks across the gene body (**Fig. 2C**). To explore how other intron features affect chromatin marks, we created a Shiny app that can be accessed at https://github.com/AlicePierce/IntronArchitecture. These observations underscore the influence of intron architecture on the distribution of active chromatin features.

### Gene duplicates with more introns have higher abundance of H3K36me3, Tissue Breadth, H2A.X, CG Methylation, and H3K4me1

To investigate the potential causal relationship between intron number and the abundance of chromatin features within gene bodies, we used gene duplicates that have diverged in intron architecture, and explored the impact of intron loss and gain on chromatin features. We profiled 9,989 pairs of *Arabidopsis* gene duplicates (Kenchanmane Raju et al. 2023). Of these, 3,786 pairs displayed differences in intron number, with an average difference of 1.88 introns per pair.

To assess the effect of intron number divergence on chromatin features, we compared the mean scaled read depth of chromatin features between gene duplicates with fewer versus more introns (**Fig. 3, Fig. S4-10**). Our analysis revealed that gene duplicates with more introns exhibit a greater abundance of chromatin features across their gene bodies that are commonly associated with active gene expression, including H3K36me3, H2A.X, CG methylation, and H3K4me1 (**Fig. 3A**). Additionally, gene duplicates with more introns have broader tissue expression and lower coefficients of variation in expression across tissues (**Fig. 3D**). Conversely, gene duplicates with fewer introns have a higher abundance of chromatin marks typically associated with repressed gene expression when found throughout the gene body, such as H2A.Z, H3.1, H3K4me2, and H3K27me3 (**Fig. 3C**).

**Figure 3.**
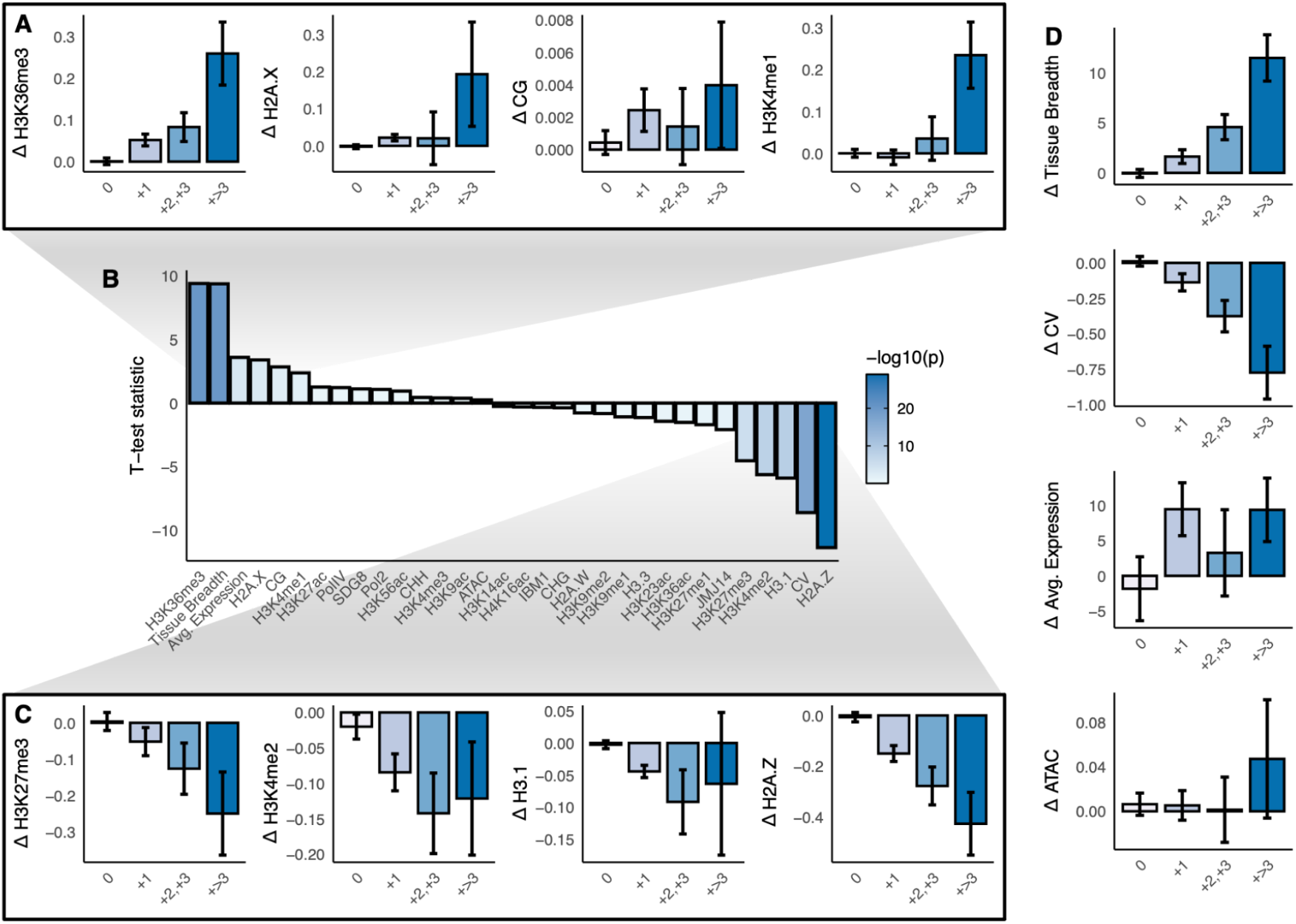
Difference in chromatin feature abundance on gene duplicates divergent in intron architecture. **(A)**Bar graphs showing the scaled read depth between gene duplicates differing in intron architecture for H3K36me3, H2A.X, CG methylation, H3K4me1. Error bars represent ± standard error. **(B)** T-test statistic between the difference in scaled read depth of duplicates with less and more number of introns in chromatin features, DNA-associated proteins, and expression between gene duplicates differing in intron architecture. **(C)** Same as (A), for H3.1, H3K27me3, H3K4me2, and H2A.Z, and **(D)** tissue breadth, coefficient of variation of expression, average expression, chromatin accessibility (ATAC-seq) N: 0 = 6203; +1 = 2461; +2,+3 = 959; +>3 = 366.

To ensure that the patterns observed are not affected by the age of gene duplication events, we categorized genes by duplication event type and found that the patterns remain consistent across different duplication events (**Fig. S5-7**). These findings highlight the potential role of intron architecture, particularly the number of introns, in shaping the chromatin landscape and regulating the gene expression of their genes.

### Transposed duplicates with more introns have higher abundance of H2A.X and CG Methylation but no change in H3K36me3

Despite the interesting results from the gene duplicate analysis, the inability to determine the ancestral state of gene duplicates limits our ability to conclude whether changes in chromatin marks are solely due to intron architecture or influenced by other factors, such as changes in promoter regions. To further assess the effects of intron loss and gain on chromatin structure, we focused on transposed gene duplicates where the parental and transposed duplicates are known (Kenchanmane Raju et al. 2023). We examined changes in chromatin mark abundance between transposed duplicates that have gained, lost, or have equal number of introns (**Fig. 4, Fig. S11-14**).

**Figure 4.**
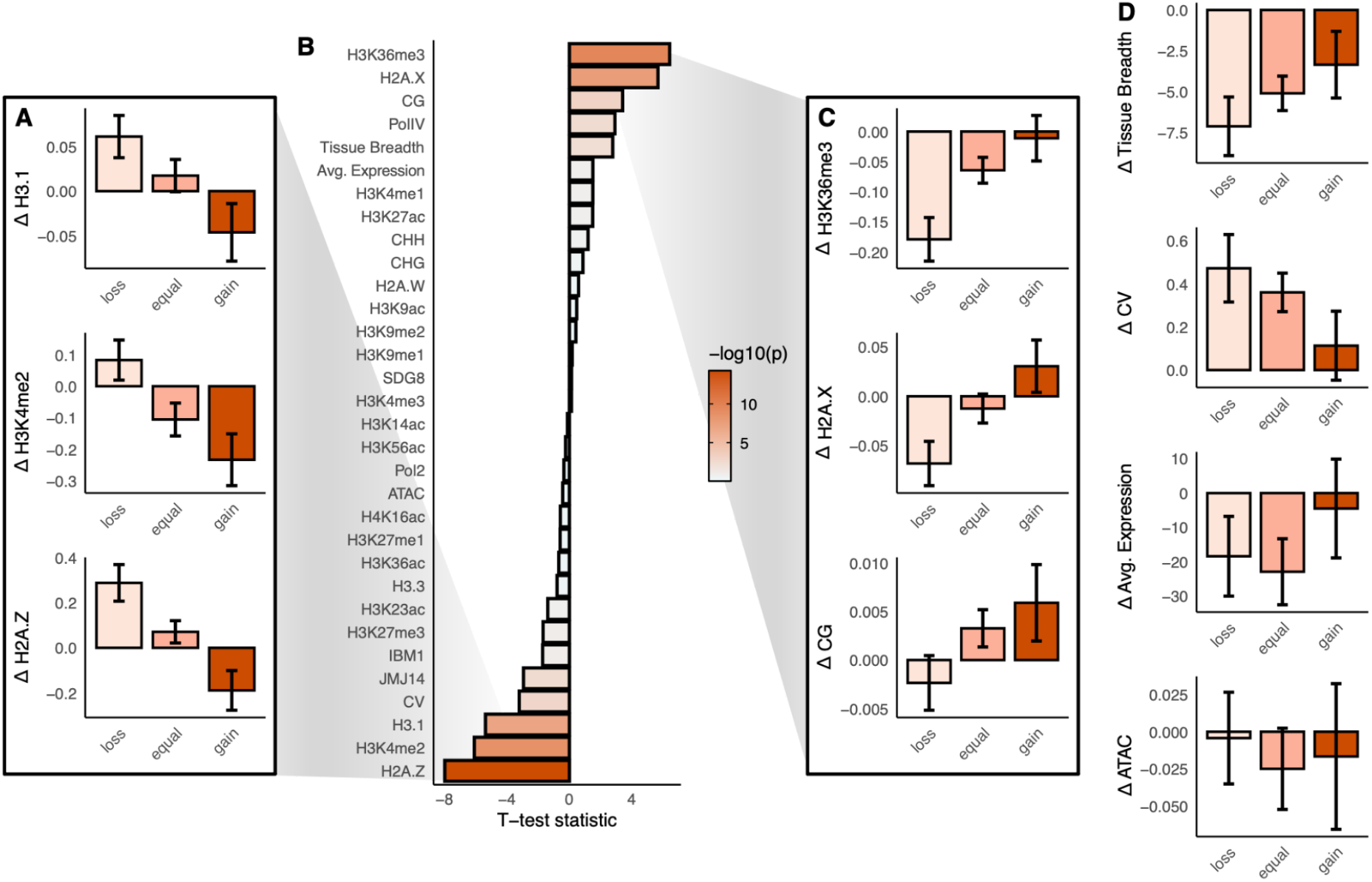
Difference in chromatin feature abundance on transposed duplicates that have lost or gained introns. **(A)**Bar graph showing the difference in scaled read depth between transposed duplicates that have lost, gained or have equal number of introns for H3.1, H3K4me2, H2A.Z. Error bars represent ± standard error. N: loss = 497; equal = 1109; gain = 356. **(B)** T-test statistic between the difference in scaled read depth of transposed duplicates that have gained or lost introns for chromatin features, DNA-associated proteins, and expression. **(C)** Same as (A) for H3K36me3, H2A.X, CG methylation **(D)** tissue breadth, coefficient of variation of expression, average expression, and chromatin accessibility (ATAC-seq)

Our analysis showed that transposed duplicates that gained introns have higher levels of H2A.X and CG methylation but lower levels of H3.1, H3K4me2, and H2A.Z (**Fig. 4AB**). Interestingly, gaining introns in transposed duplicates did not appear to affect the abundance of H3K36me3 (**Fig. 4B**), suggesting that intron gain is accompanied by changes in chromatin states that are largely independent of gene body histone marks linked to transcriptional regulation.

In contrast, transposed duplicates that lost introns exhibit a higher abundance of H3.1, H3K4me2, and H2A.Z (**Fig. 4A**), and have decreased abundance of H3K36me3, H2A.X, and CG methylation (**Fig. 4B**). These duplicates also demonstrated lower tissue breadth and higher coefficients of variation of expression across tissues (**Fig. 4D**). These findings provide further evidence of the association of intron architecture on chromatin features and gene expression regulation, and highlight the distinct chromatin states and expression patterns associated with intron loss and gain in transposed duplicates.

## Discussion

Here we present evidence supporting that intron architecture predicts the chromatin landscape of their associated genes. We observe a positive correlation between first intron position and active histone marks commonly enriched near transcription start sites, such as H3K4me3, H3K4me2, H3K23ac, and H3K14ac (Zhang et al. 2009; Leng et al. 2020; van Dijk et al. 2010; Stroud et al. 2014; Chen et al. 2017). This observation supports previous hypotheses that first introns may facilitate the deposition of histone post-translational modifications near transcription start sites, which might positively regulate transcription (Gallegos and Rose 2015). However, our findings contrast work that demonstrated that stimulatory intronic sequences closer to the transcription start site enhance gene expression (Rose 2004). Instead, we found that the position of the first intron is positively correlated with both transcription start site-associated histone marks and gene expression, suggesting a different general regulatory role of the first intron position relative to the transcription start site, versus the position of stimulatory sequences found within certain first introns.

Interestingly, our results also show that the amplitude of histone marks commonly enriched near transcription start sites, such as H3K4me3, H3K4me2, H3K23ac, and H3K14ac, increases with greater first intron distance from the transcription start site. This finding diverges from observations in human cell lines, where a closer first intron position correlates with a higher peak of H3K4me3 (Bieberstein et al. 2012). Contrary to expectations, we did not observe a peak shift upstream relative to the TSS, regardless of intron position. In contrast, human cell lines display a peak shift and a second promoter-proximal peak when the first intron is further from the TSS. This suggests that the influence of first intron position on the deposition of chromatin marks near the TSS has diverged between animals and plants, which could indicate different intronic regulatory mechanisms at play.

Our analysis also reveals that intron number is associated with active chromatin features commonly distributed throughout gene bodies, such as H3K4me1, H2A.X, and H3K36me3. In plants, H3K4me1 is associated with actively transcribed regions and DNA repair (Niu et al. 2021; Quiroz et al. 2024; Zhang et al. 2009; Lu et al. 2019). H2A.X has been implicated in DNA double-strand break response (Roitinger et al. 2015; Lang et al. 2012; Charbonnel et al. 2011), as well as transcriptional activation (Xiao et al. 2021). H3K36me3 is hypothesized to have a role in RNA Polymerase elongation and stalling (Kindgren et al. 2020; Leng et al. 2020). Our results are supported by recent findings that suggest that different H3K36 methyltransferases act on different genes depending on their gene structure (Yao et al. 2025). For instance, SDG8, believed to be the major H3K36 methyltransferase in *Arabidopsis*, was found to suppress the H3K36 methyltransferase activity of SDG4 on short and intronless genes. However, SDG8 works together with SDG25 and SDG26 to deposit H3K36me3 on longer genes with a higher number of introns (Yao et al. 2025). Further investigation into whether the number of introns in the gene body plays a role in determining the abundance of these chromatin features in gene bodies, and their effect on expression, will be important for understanding the interplay between intron architecture and the epigenome.

We sought to determine whether chromatin features associated with intron architecture exhibit differential distribution between introns and exons. In humans, H3K36me3 is enriched in exons relative to introns (Frigola et al. 2017), with similar patterns observed in *C. elegans* and fission yeast (Wilhelm et al. 2011; Kolasinska-Zwierz et al. 2009; Tilgner et al. 2009). In contrast, in *Arabidopsis*, H3K36me3 displays a positional gradient distribution along ordinal exons and introns, with its abundance increasing in genes containing a higher number of introns. This suggests that plants may employ distinct mechanisms for H3K36me3 deposition and maintenance along the gene body. Other marks, such as H3K4me1, also exhibit enrichment patterns that vary with ordinal exon and intron position along the gene body. This is consistent with the idea that H3K4me1 acts as a transcriptional “memory” mark, reflecting recent transcriptional activity (Oya et al. 2022; Krogan et al. 2003; Ng et al. 2003; Soares et al. 2017; Bae et al. 2020).

In humans, H3K36me3 is linked to the recruitment of mismatch repair mechanisms at actively transcribed genes to prevent mutations (Huang et al. 2018; Girardini et al. 2023). H3K36me3 is recognized by the MSH6 DNA repair protein through its PWWP domain (Li et al. 2013; Sun et al. 2020). However, in *Arabidopsis*, MSH6 lacks the PWWP domain and instead contains a Tudor domain that recognizes H3K4me1 (Quiroz et al. 2024). This divergence in domain structure may partially explain the differential distribution of H3K36me3 in *Arabidopsis*, possibly facilitating a shift in its distribution in gene bodies. Beyond its role in mismatch repair, H3K36me3 has also been implicated in the regulation of alternative splicing in both plants and humans (Sharda and Humphrey 2022; Pradeepa et al. 2012; Guo et al. 2014; Luco et al. 2010; Andersson et al. 2009; Luco et al. 2011; Kolasinska-Zwierz et al. 2009; Leng et al. 2020; Pajoro et al. 2017; Wei et al. 2018), suggesting a broader functional impact on gene expression beyond chromatin association.

To further investigate whether introns directly influence chromatin feature distribution, we analyzed the effect of intron divergence in gene duplicates, including those arising from transposition events. Our results corroborated the findings from the global analysis, showing that gene duplicates with a higher number of introns are enriched for chromatin features positively associated with gene expression, such as H3K36me3, H2A.X, CG methylation, and H3K4me1, as well as tissue breadth.

Moreover, we observed that transposed duplicates that have gained introns are enriched for H2A.X and CG methylation. Notably, intron gain in transposed duplicates did not increase H3K36me3 abundance, which may be explained in part by the general trend of transposed genes having lower expression compared to their source copy, and therefore having a decrease on activating histone marks. Indeed, transposed genes that lost introns showed a decrease in H3K36me3, beyond that seen in transposed genes with no change in intron number. When examining the effects of transposition on chromatin marks, we found that gaining introns often reversed or mitigated the abundance of chromatin marks associated with repressed gene expression when found throughout the gene body, such as H3.1, H3K4me2, and H2A.Z (Coleman-Derr and Zilberman 2012; Sura et al. 2017; Stroud et al. 2012; Wollmann et al. 2012; Foroozani et al. 2022; Cortijo et al. 2017; Liu et al. 2019). This leads us to hypothesize that introns may play a role in distinguishing functional genes from transposable elements or pseudogenes that arose from retrotransposition.

While we acknowledge that the protein-coding sequence also diverges in gene duplicate events, our null hypothesis is that no association should exist between intron features and chromatin features—yet we consistently observe such associations. Additionally, previous literature indicates that gene expression changes are frequently driven by non-coding sequences rather than protein-coding sequences. In tissue-specific gene duplicates, however, we cannot exclude the possibility that the chromatin state may differ in tissues where these genes are predominantly expressed, particularly since the ChIP-seq data we accessed were derived from leaf tissue, which may not fully represent chromatin dynamics in other tissue contexts.

## Conclusions

Our findings provide new insights into how introns regulate chromatin biology, emphasizing their potential roles in transcriptional regulation. By identifying correlations between intron architecture and chromatin features, we found that the first intron position can predict chromatin marks that are found near the TSS, such as H3K4me3, H3K4me2, H3K23ac, and H3K14ac. We also found that more introns in genes are associated with the abundance of active gene body marks such as H3K4me1, and H3K36me3. These results suggest that introns influence gene activity by modulating the chromatin state of their genes in potentially multiple ways.

Notably, we found that the chromatin mark H3K36me3 is distributed across both exons and introns in *Arabidopsis*, with its abundance increasing with intron number and varying with position along the gene body. This contrasts with previous findings in humans, *C. elegans*, and yeast, where H3K36me3 is enriched in exons, suggesting a divergence in the role of H3K36me3 on gene regulation between plants and other eukaryotes.

We propose two mechanisms through which introns might promote gene expression in their associated genes:

1. First introns promote transcription initiation:
  - First introns promote the accumulation of histone marks near the transcription start site, facilitating transcription initiation.
2. More introns in the gene body promote histone mark accumulation:
  - An increased number of introns could extend gene body length, providing more space for histone marks to accumulate, thereby promoting gene expression.
  - Canonical intron-exon junction motifs, or other intronic sequences, may be recognized by histone writers, with more introns facilitating more recognition spots that allow the accumulation of gene body marks associated with active transcription.

The patterns revealed in this work, and the proposed mechanisms underlying them, emphasize the potential regulatory roles of introns in shaping the chromatin state and the transcriptional state of their genes, warranting further experimental investigation to clarify their functions in gene expression and genome biology.

## Acknowledgments

We would like to thank members of the Monroe lab for their support and discussions about the

## Supplemental Materials

This work was supported by National Science Foundation Grants 2317191 and 2338236 to J.G.M.. Research was conducted at the University of California Davis, which is located on land that has been the home of the Patwin people for thousands of years.

## Code and data availability

Code for this research is maintained at https://github.com/AlicePierce/IntronArchitecture. This repository contains scripts to analyze the data and create figures. In addition, it contains information on how to download and run Shiny apps to further explore the association between intron architecture and chromatin features in *Arabidopsis*.

## Author Contributions

Conceptualization: AP, AR, GM

Methodology: AP, GM

Software: AP

Validation: AP, GM

Formal Analysis: AP, GM

Data Curation: AP, GM

Writing - Original Draft: AP

Writing - Review & Editing: AP, AR, GM

Visualization: AP, GM

Supervision: GM, AR

Funding Acquisition: GM

## Supplemental tables

**Supplementary Table S1**. ChIP-seq and ATAC-seq file correlations

**Supplementary Table S2**. Pairwise Spearman Correlation Rho between intron architecture and chromatin features

**Supplementary Table S3**. Statistical Significance (p-values) of Spearman Correlations between intron architecture and chromatin features

**Supplementary Table S4**. Random Forest Model Performance Metrics

**Supplementary Table S5**. Random Forest Model Feature Importance

## Supplemental figures

**Figure S1.**
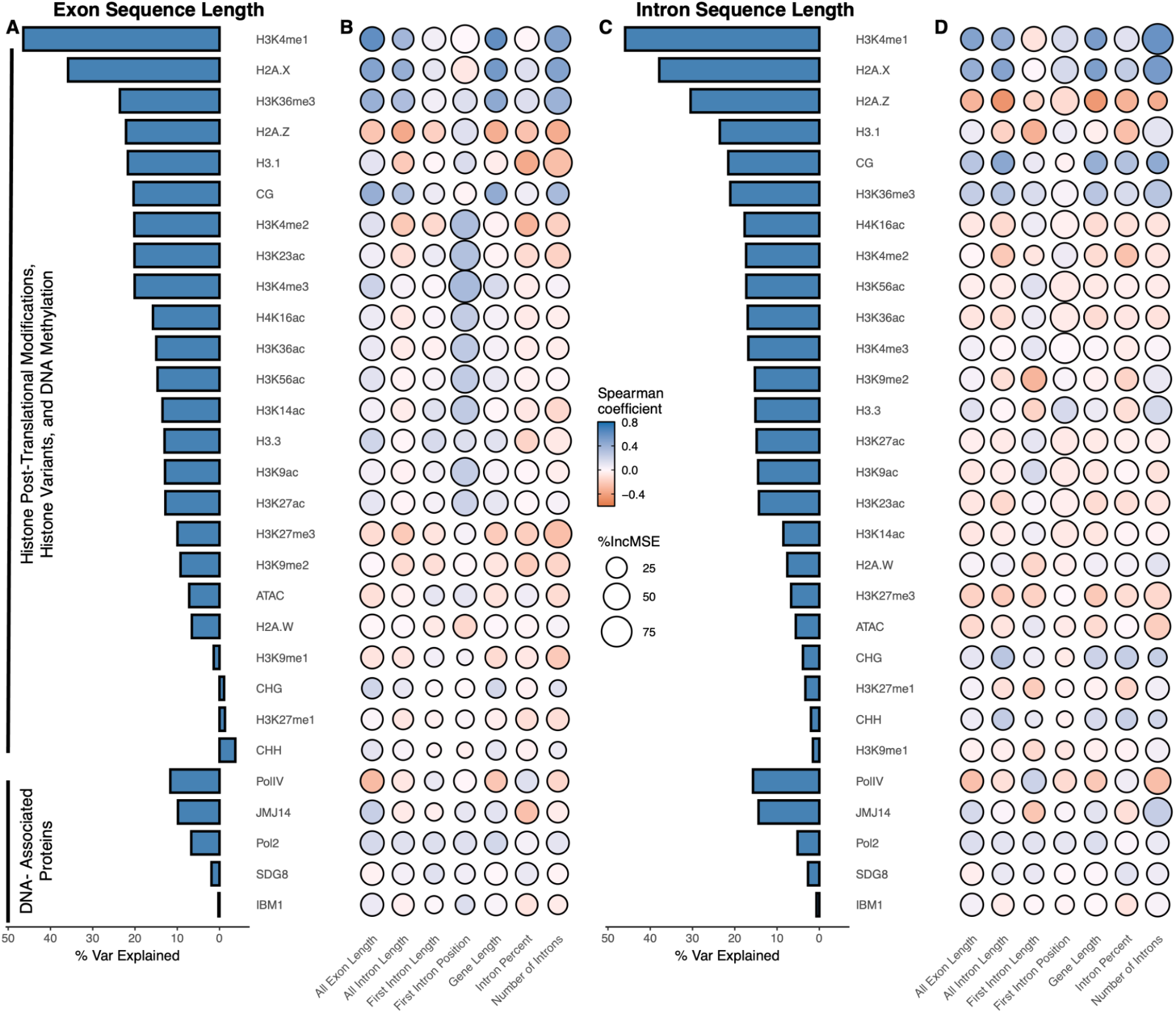
Prediction of chromatin features from intron architecture. **(A)** Bar graph showing the percent variance explained from random forest regression between gene features predicting each chromatin feature, DNA-associated protein or gene expression feature across exon sequence length and **(B)** Correlation heatmap between gene architecture, and chromatin features, DNA-associated features or gene expression features across exon sequence length. **(C)** Same as (A) across intron sequence length **(D)** Same as (B) across intron sequence length. The color indicates the Spearman correlation coefficient between gene features and each chromatin feature. Size indicates the percent increase in mean square error introduced in a random forest regression when the corresponding predictor is permuted.

**Figure S2.**
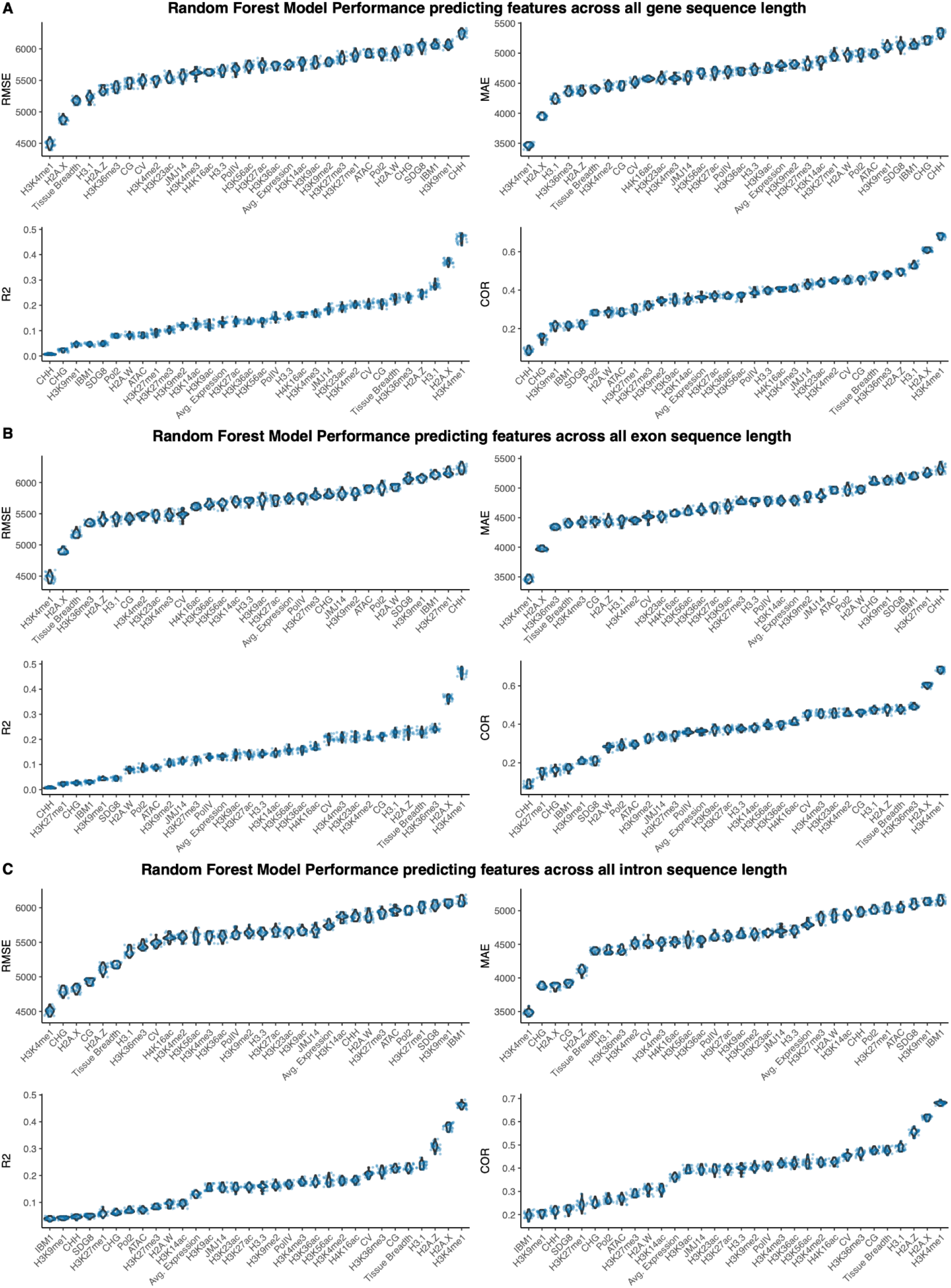
Model Performance Metrics for Random Forest Predictions. Violin plots with superimposed boxplots depicting the RMSE, MAE, R2, and Pearson correlation between actual and predicted values across **(A)** all gene sequence length **(B)** exon sequence length **(C)** intron sequence length

**Figure S3.**
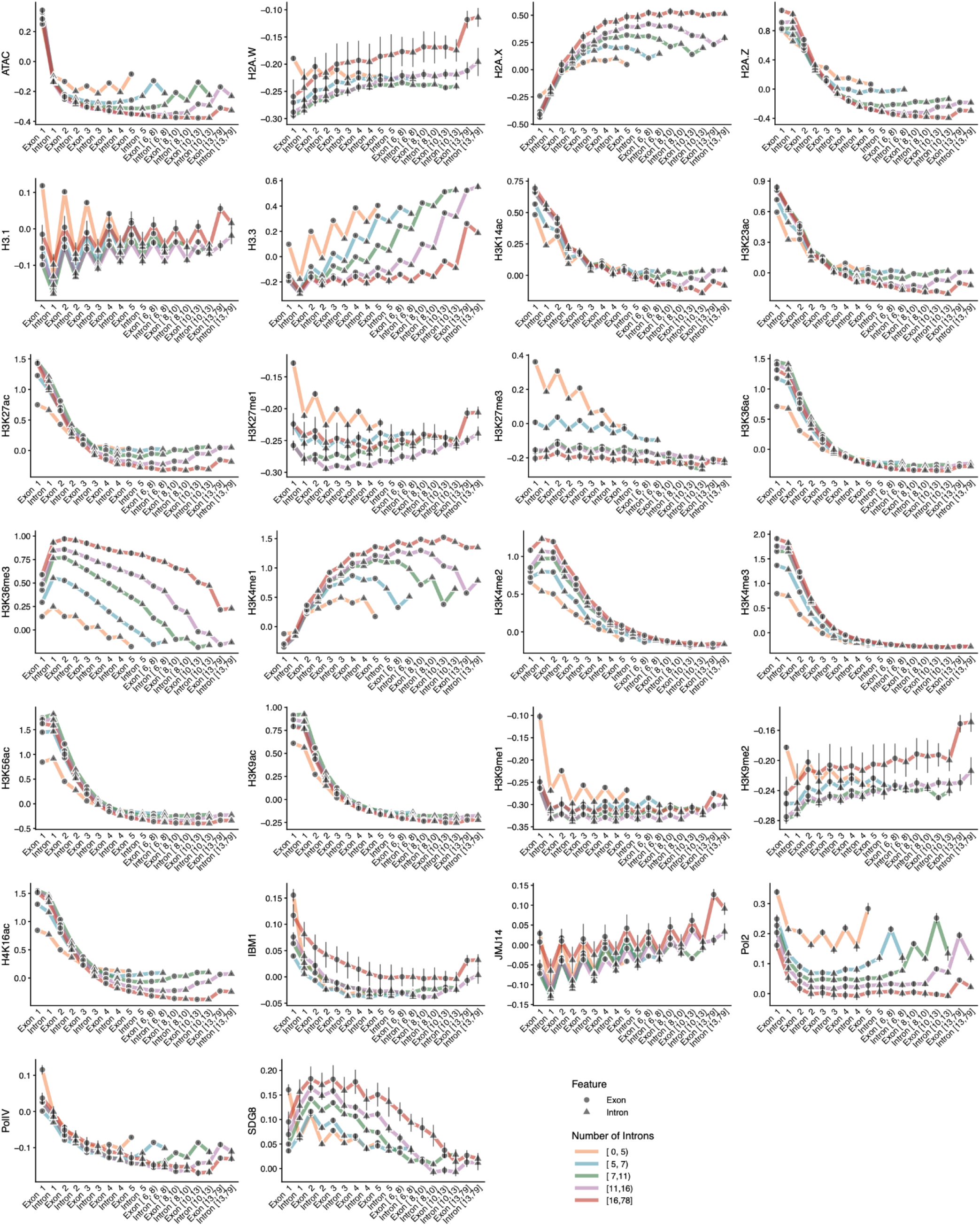
Scatterplots of chromatin features across ordinal number of exons and introns. Each plot shows the mean abundance of chromatin marks across ordinal exons or introns. Genes were binned according to their number of introns. Circles represent exons and triangles represent introns. Error bars represent ± standard error.

**Figure S4.**
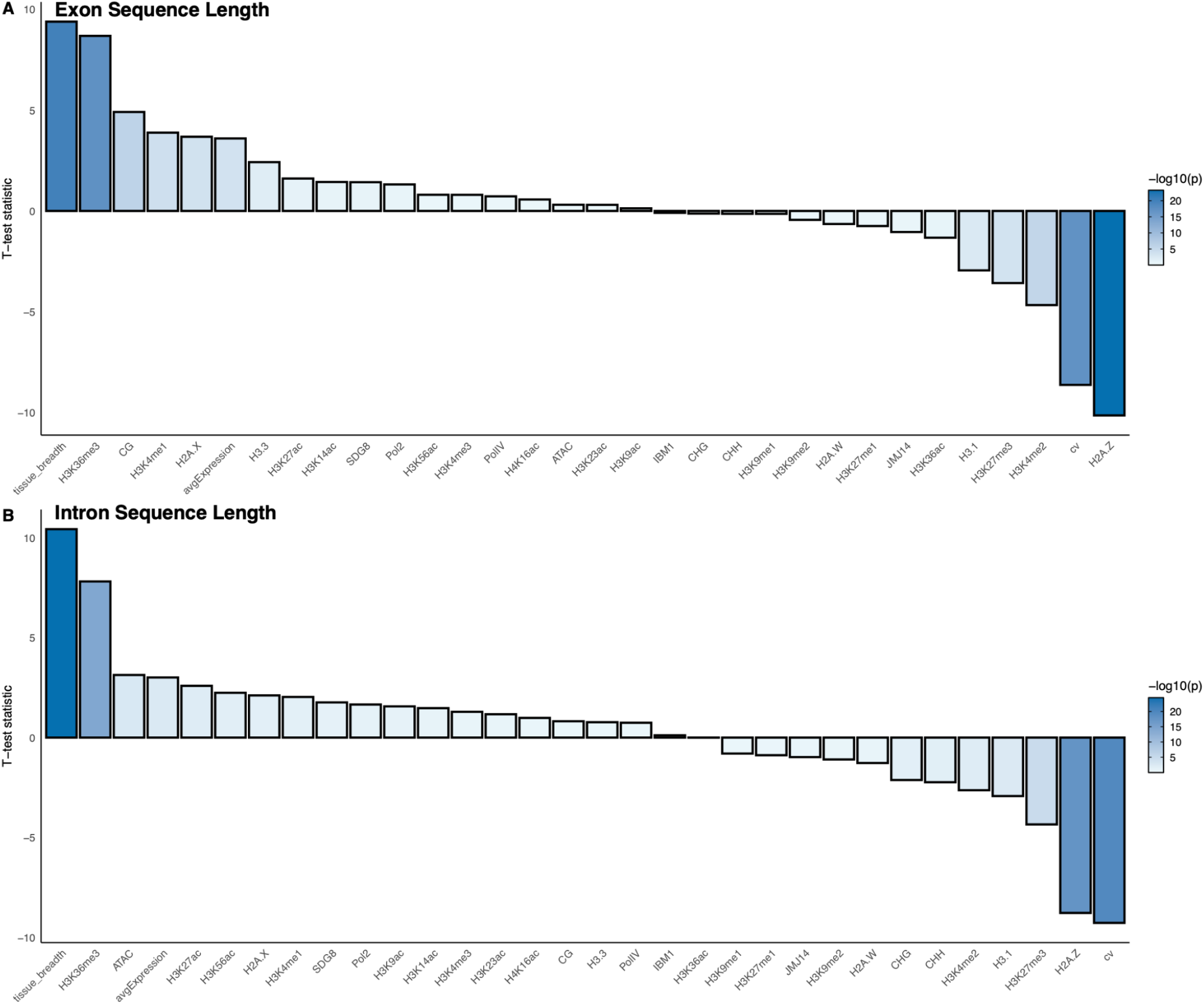
Difference in chromatin feature abundance on gene duplicates. T-test statistic between the difference in scaled read depth of gene duplicates with less and more number of introns across **(A)** exon sequence length **(B)** intron sequence length.

**Figure S5.**
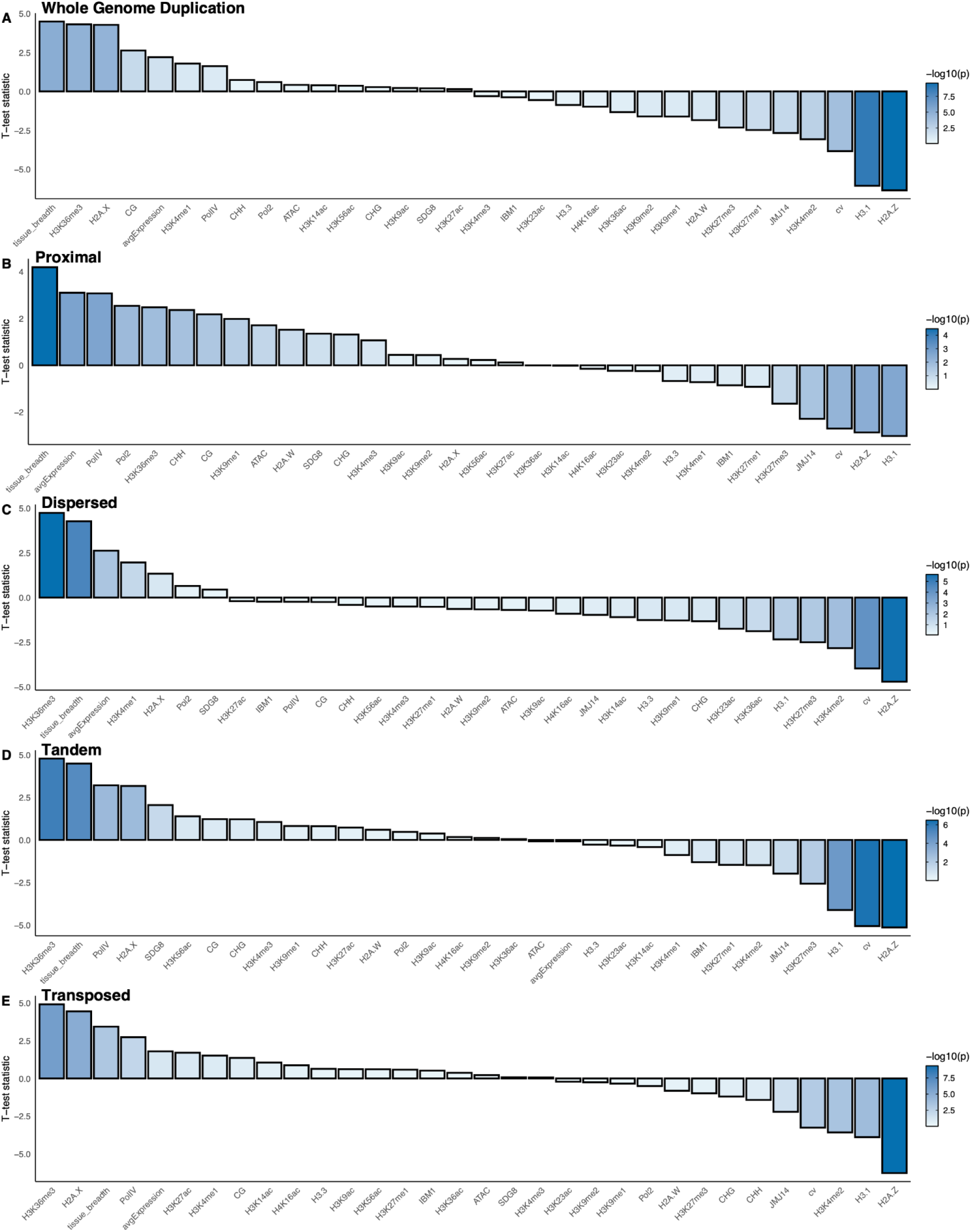
Difference in chromatin feature abundance on gene duplicates across gene duplication events across gene sequence length. T-test statistic between the difference in scaled read depth of gene duplicates with less and more number of introns across gene sequence length, binned by the following gene duplication events: **(A)** Whole Genome Duplication **(B)** Proximal **(C)** Dispersed **(D)** Tandem **(E)** Transposed.

**Figure S6.**
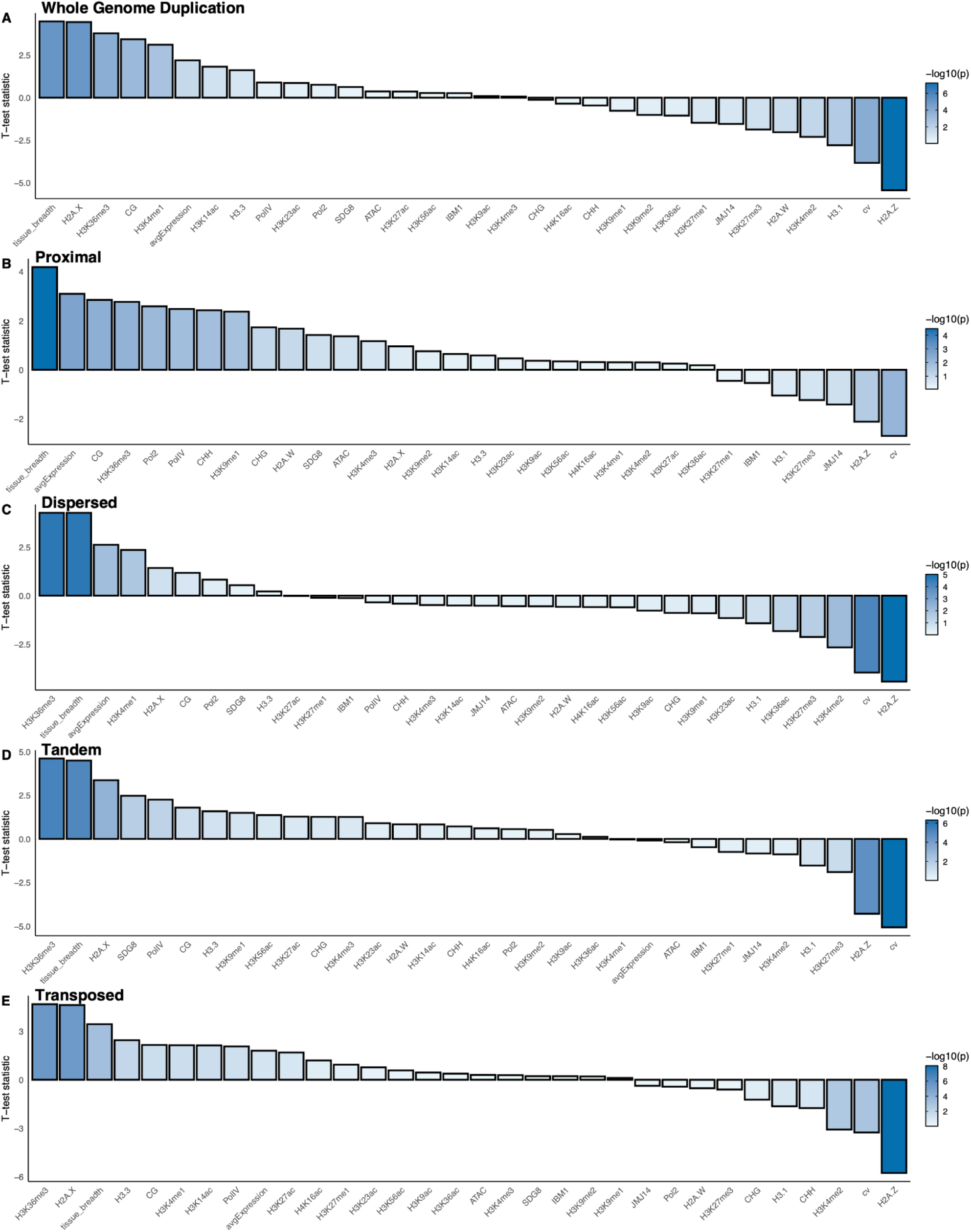
Difference in chromatin feature abundance on gene duplicates across gene duplication events across exon sequence length. T-test statistic between the difference in scaled read depth of gene duplicates with less and more number of introns across exon sequence length, binned by the following gene duplication events: **(A)** Whole Genome Duplication **(B)** Proximal **(C)** Dispersed **(D)** Tandem **(E)** Transposed.

**Figure S7.**
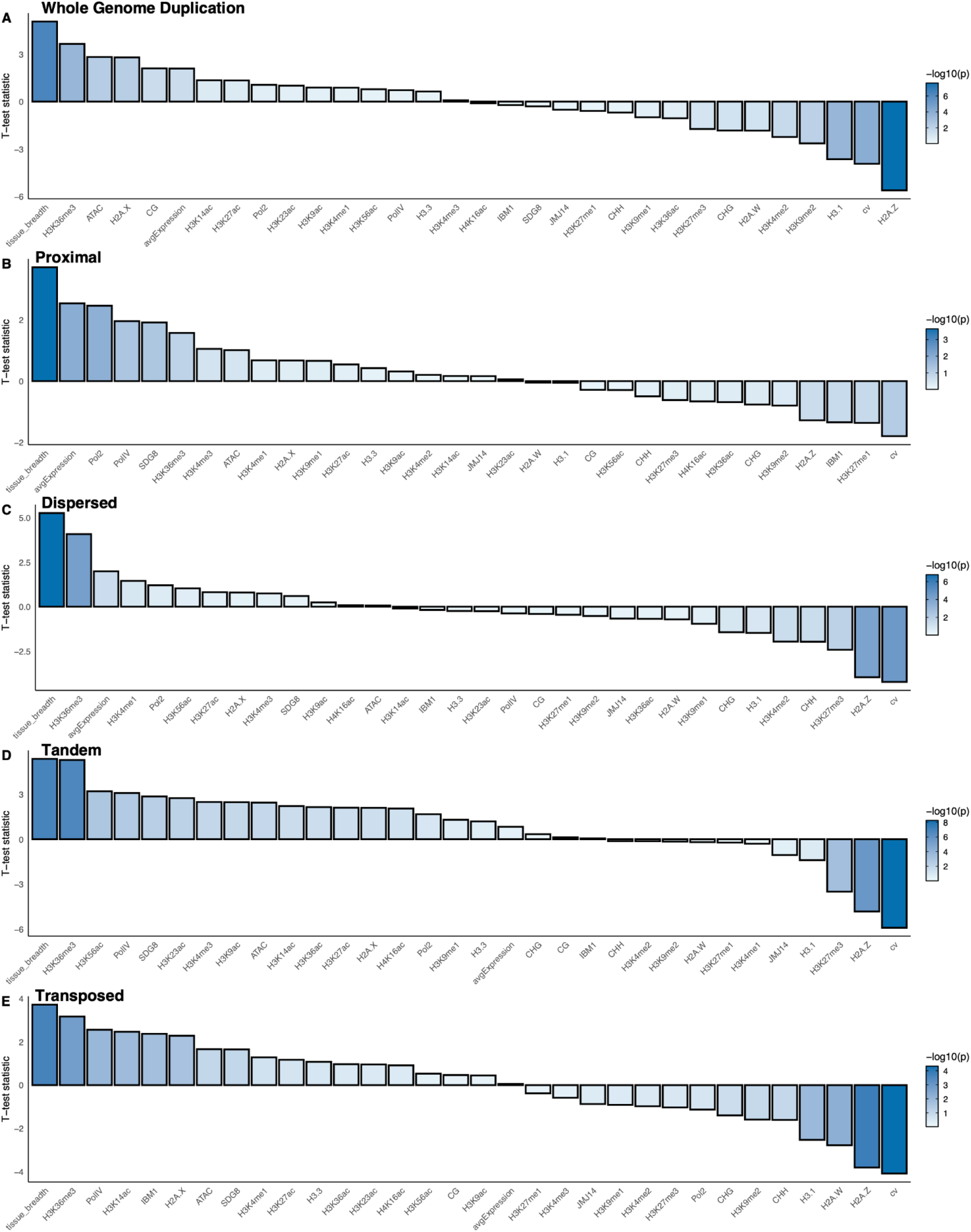
Difference in chromatin feature abundance on gene duplicates across gene duplication events across intron sequence length. T-test statistic between the difference in scaled read depth of gene duplicates with less and more number of introns across intron sequence length, binned by the following gene duplication events: **(A)** Whole Genome Duplication **(B)** Proximal **(C)** Dispersed **(D)** Tandem **(E)** Transposed.

**Figure S8.**
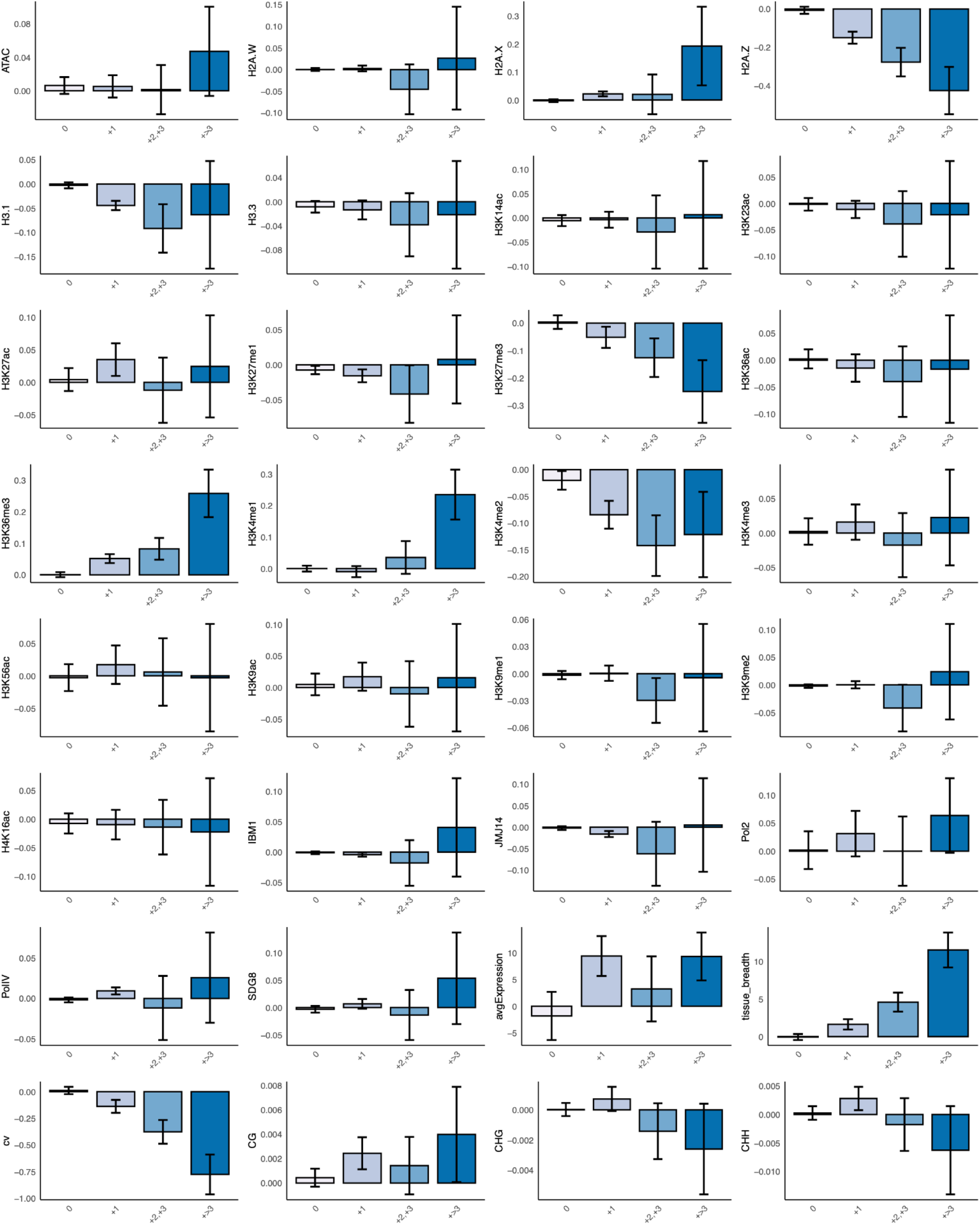
Difference in chromatin feature abundance across gene duplicates with divergent intron architecture across gene sequence length. Bar graphs showing the difference in scaled read depth across gene duplicates that differ in intron number across gene sequence length. Error bars represent ± standard error. N: 0 = 6203; +1 = 2461; +2,+3 = 959; +>3 = 366.

**Figure S9.**
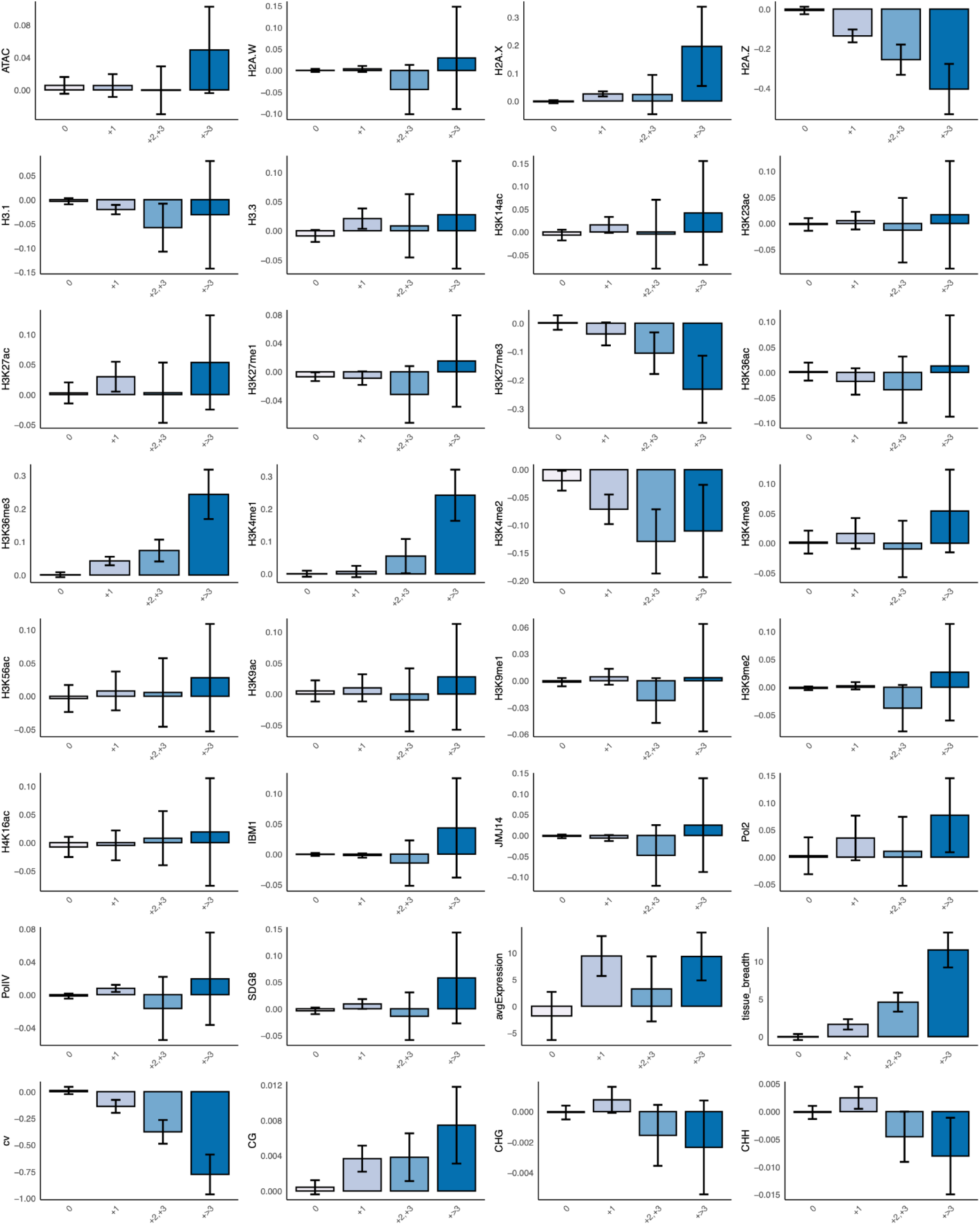
Difference in chromatin feature abundance across gene duplicates with divergent intron architecture across exon sequence length. Bar graphs showing the difference in scaled read depth across gene duplicates that differ in intron number across exon sequence length. Error bars represent ± standard error. N: 0 = 6203; +1 = 2461; +2,+3 = 959; +>3 = 366.

**Figure S10.**
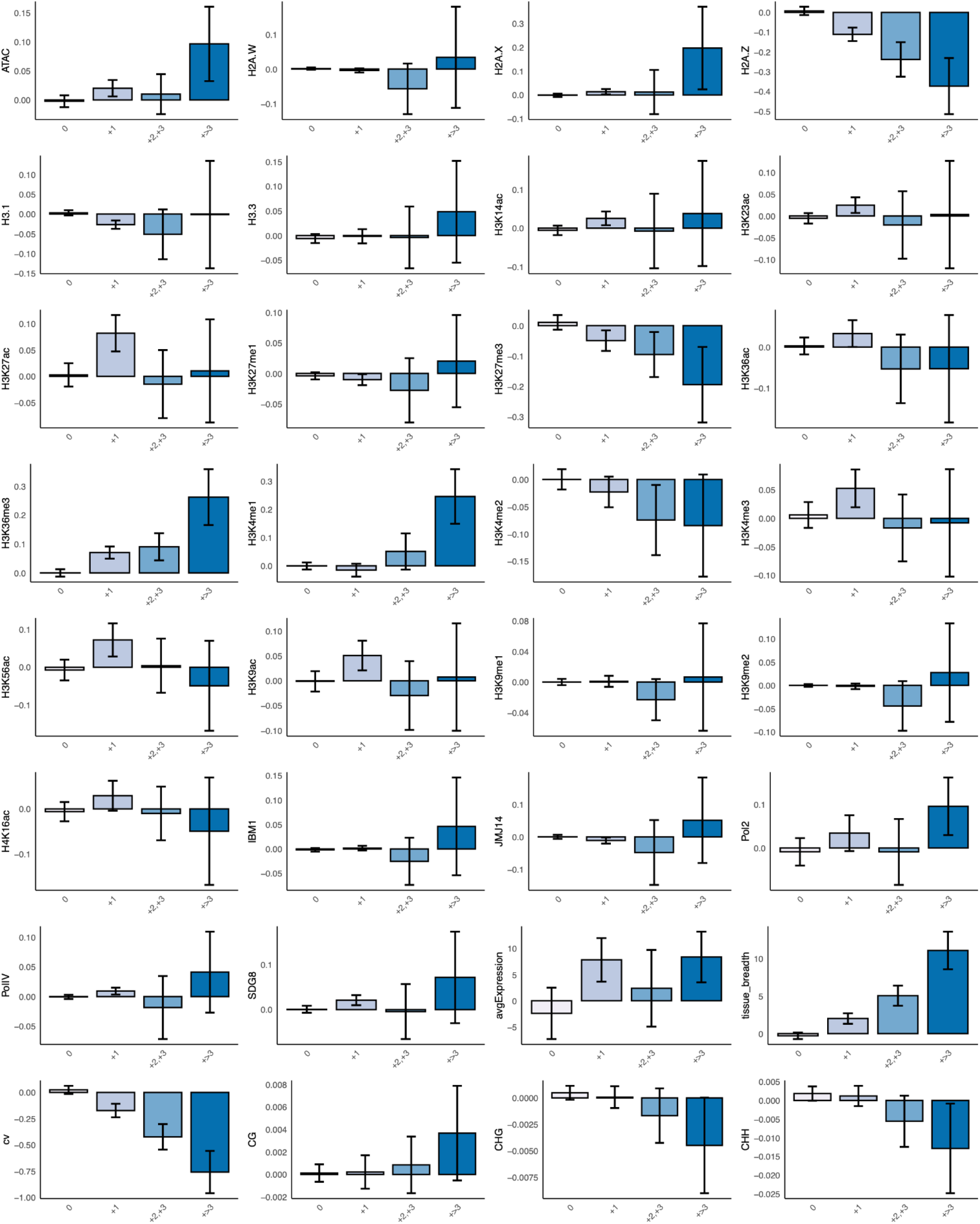
Difference in chromatin feature abundance across gene duplicates with divergent intron architecture across intron sequence length. Bar graphs showing the difference in scaled read depth across gene duplicates that differ in intron number across all intron sequence length of intron-containing genes. Error bars represent ± standard error. N: 0 = 43722; +1 = 1943; +2,+3 = 760; +>3 = 294.

**Figure S11.**
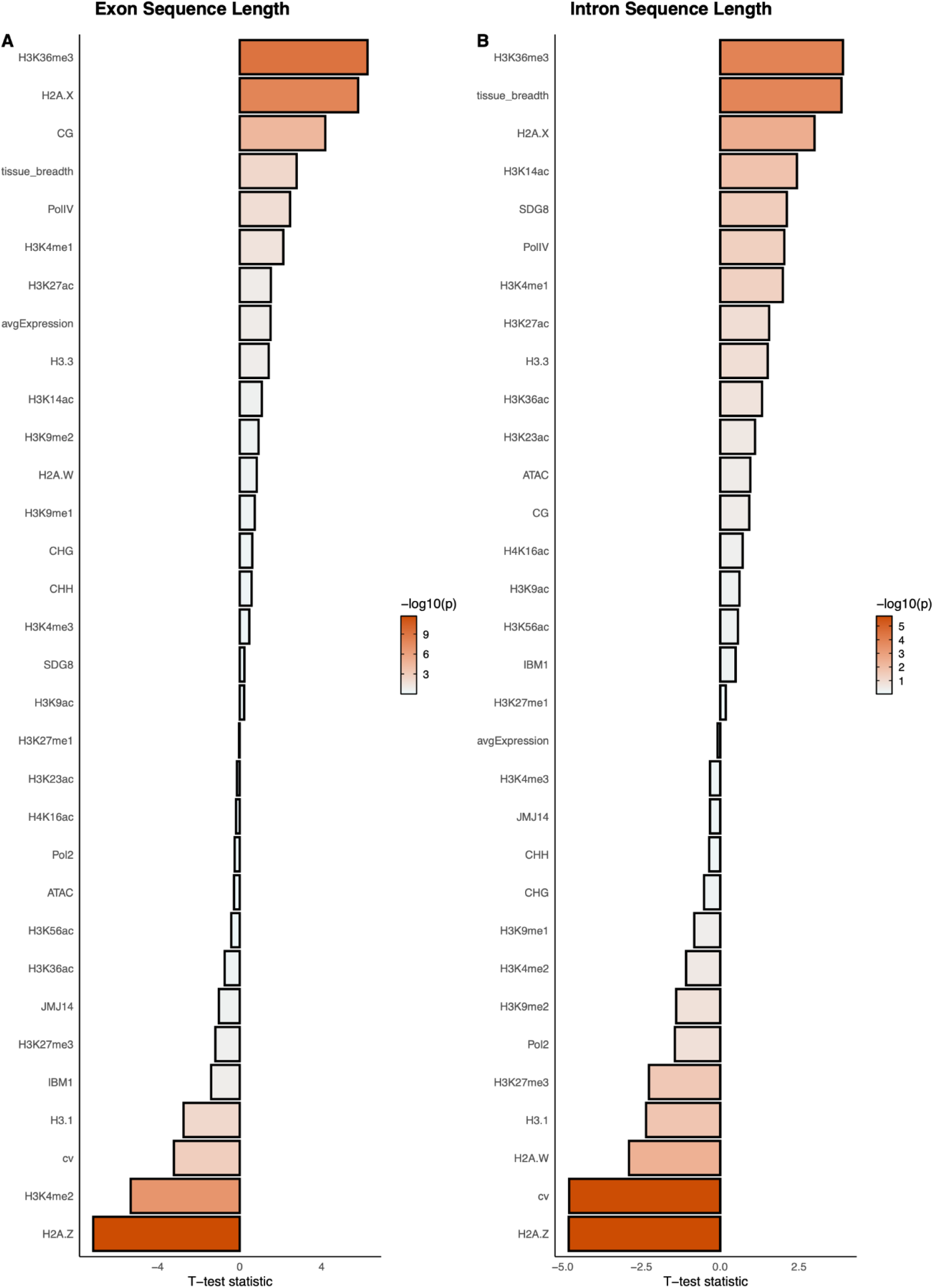
Difference in chromatin feature abundance on transposed duplicates. T-test statistic between the difference in scaled read depth of transposed duplicates that have gained or lost introns across **(A)** exon sequence length **(C)** intron sequence length.

**Figure S12.**
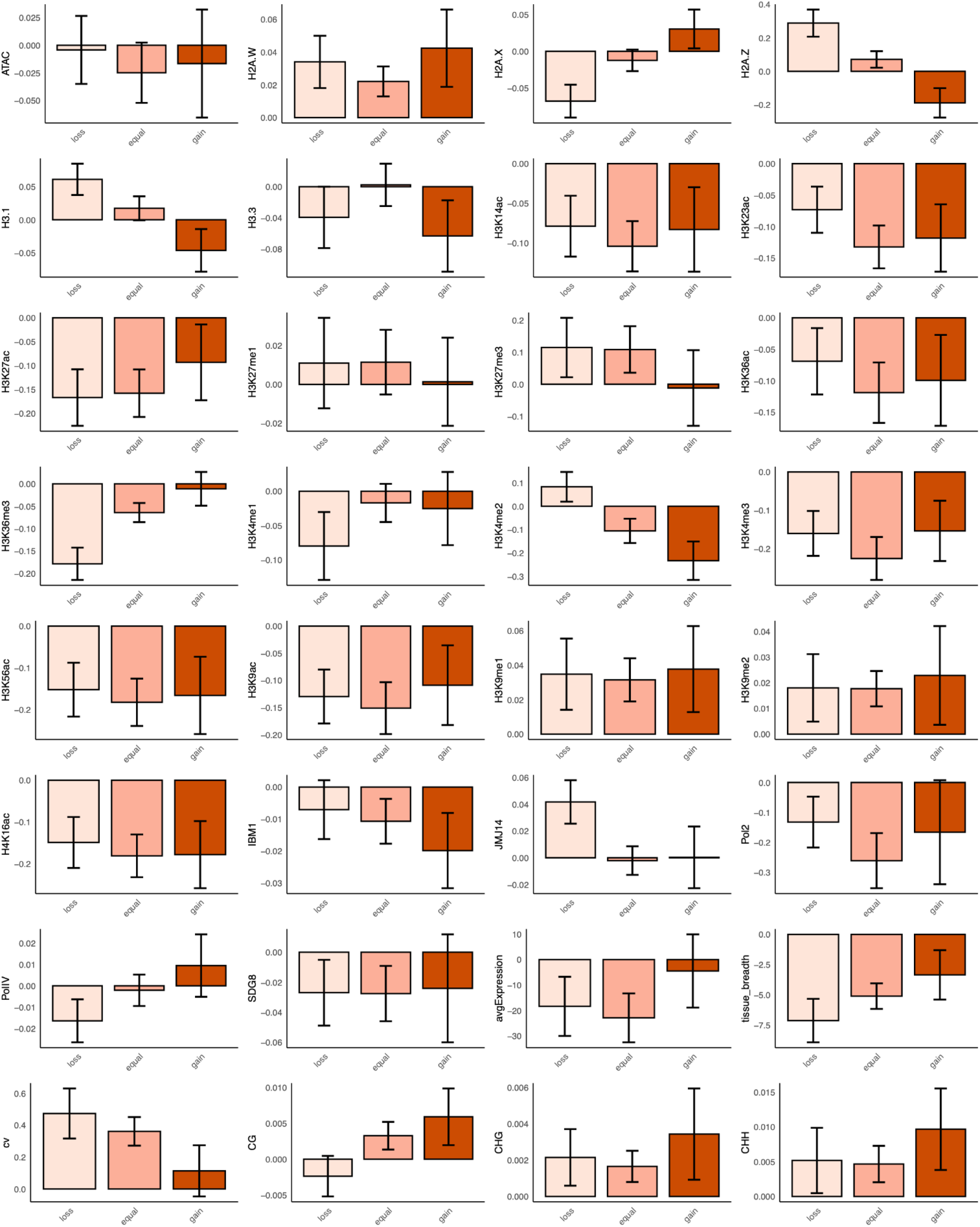
Difference in chromatin feature abundance across transposed duplicates that have lost, gained, or have equal number of introns across gene sequence length. Bar graphs showing the difference in scaled read depth across transposed duplicates that have lost, gained, or have equal number of introns across gene sequence length. Error bars represent ± standard error. N: loss = 497; equal = 1109; gain = 356.

**Figure S13.**
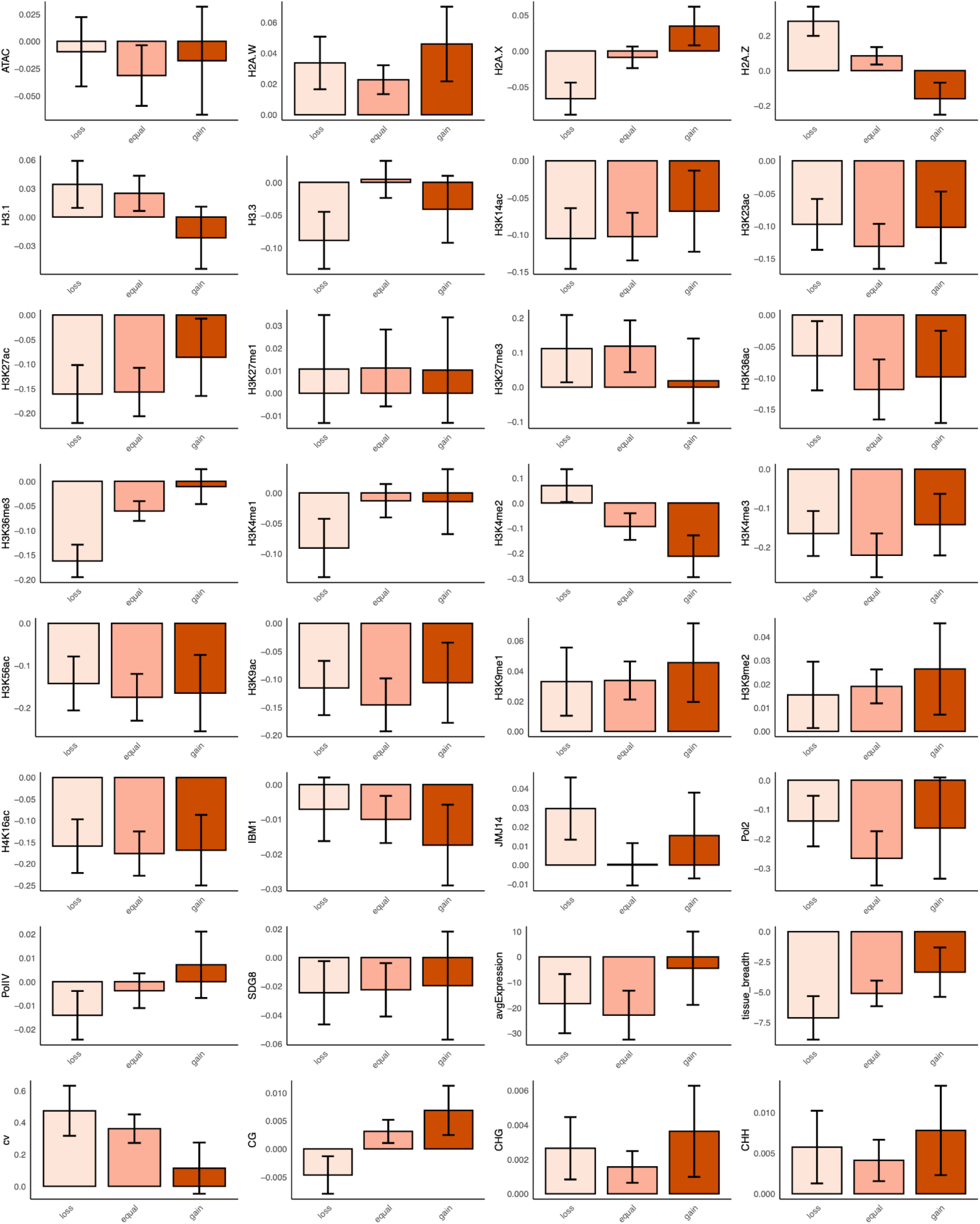
Difference in chromatin feature abundance across transposed duplicates that have lost, gained, or have equal number of Introns across exon sequence length. Bar graphs showing the difference in scaled read depth across transposed duplicates that have lost, gained, or have equal number of introns across exon sequence length. Error bars represent ± standard error. N: loss = 497; equal = 1109; gain = 356.

**Figure S14.**
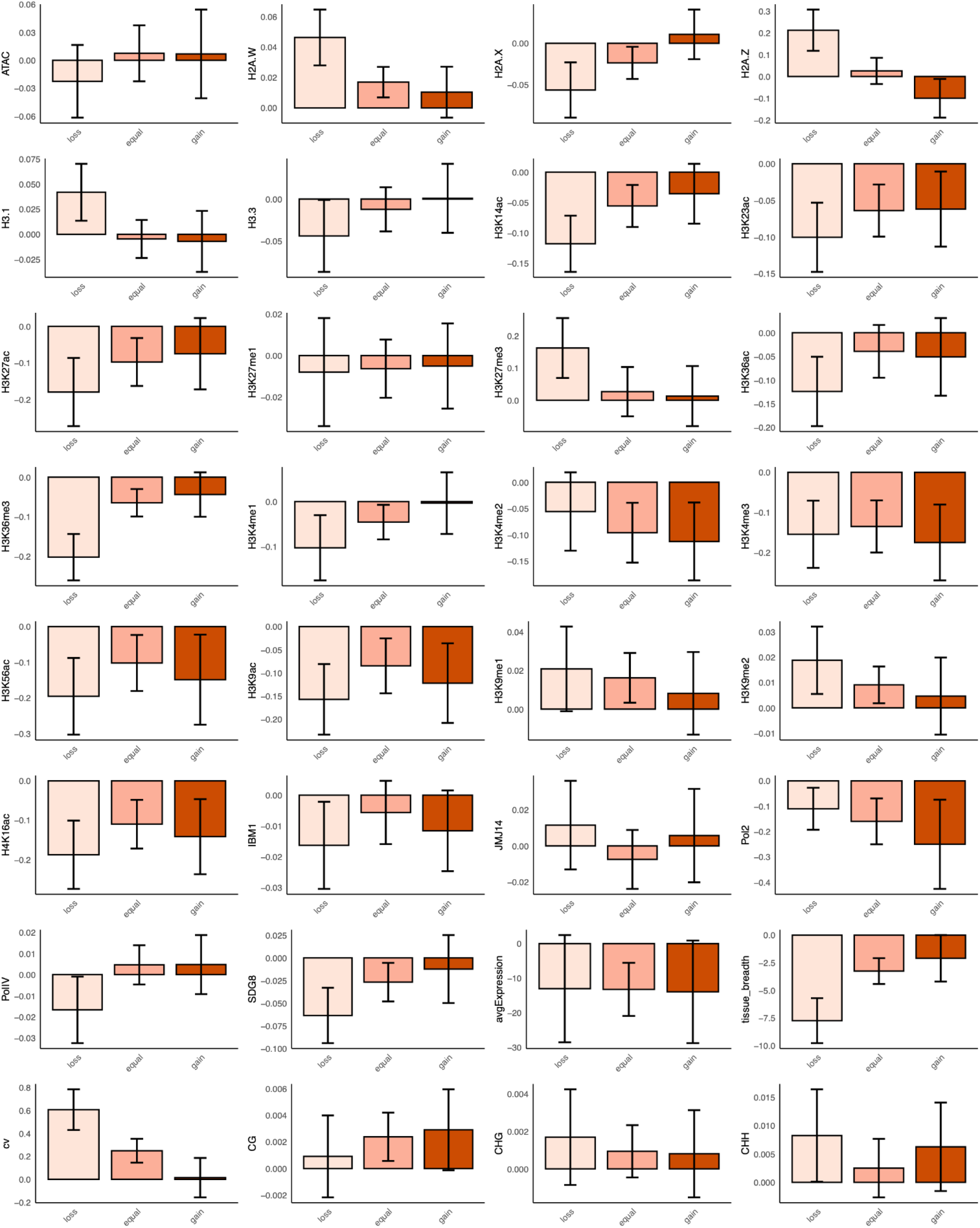
Difference in chromatin feature abundance across transposed duplicates that have lost, gained, or have equal number of introns across intron sequence length. Bar graphs showing the difference in scaled read depth across transposed duplicates that have lost, gained, or have equal number of introns across intron sequence length for intron-containing genes. Error bars represent ± standard error. N: loss = 334; equal = 735; gain = 271

